# Dynamic tripartite construct of interregional engram circuits underlies forgetting of extinction memory

**DOI:** 10.1101/2022.04.30.490143

**Authors:** Xue Gu, Yan-Jiao Wu, Jia-Jie Zhu, Xin-Rong Wu, Qi Wang, Xin Yi, Ze-Jie Lin, Zhi-Han Jiao, Miao Xu, Qin Jiang, Ying Li, Nan-Jie Xu, Michael Xi Zhu, Lu-Yang Wang, Tian-Le Xu, Wei-Guang Li

**Author notes:** Correspondance (T.-L.X.), (W.-G.L.). These authors contributed equally (Xue Gu, Yan-Jiao Wu).

## Abstract

Fear extinction allows for adaptive control of learned fear responses but often fails, resulting in a renewal or spontaneous recovery of the extinguished fear, i.e., forgetting of the extinction memory readily occurs. Using an activity-dependent neuronal labeling strategy, we demonstrate that engram neurons for fear extinction memory are dynamically positioned in the medial prefrontal cortex (mPFC), basolateral amygdala (BLA), and ventral hippocampus (vHPC), which constitute an engram construct in the term of directional engram synaptic connectivity from the BLA or vHPC to mPFC, but not that in the opposite direction, for retrieval of extinction memory. Fear renewal or spontaneous recovery switches the extinction engram construct from an accessible to inaccessible state, whereas additional extinction learning or optogenetic induction of long-term potentiation restores the directional engram connectivity and prevents the return of fear. Thus, the plasticity of engram construct underlies forgetting of extinction memory.

## Introduction

Extinction is a behavioral repertoire of the reduction of responses that occur in associative learning when reinforcers or unconditioned stimuli (US) are no longer present and thus allows for an adjustment in response to a learned behavior in a particular environment and serves as an effective treatment strategy for behavioral disorders. Auditory fear conditioning, in which the conditioned stimulus (CS; an auditory cue tone) is associated with a US, such as an electric shock, represents a powerful behavioral model used to study fear memory [1–5]. Although conditioned fear responses (i.e., freezing in rodents) can be elicited upon re-exposure to the CS, extinction via repetitive tone presentation to the conditioned subject without foot shock renders the CS memory retrieval no longer sufficient to elicit a conditioned fear response. Although the behavioral efficacy of extinction for suppressing fear memory expression is encouraging, it is more important to note its inherent instability [6–8]. Indeed, extinguished fear can be readily renewed when subjects are presented with CS in any context other than that in which extinction learning was performed, a process termed as fear renewal [9–11], or extinguished fear can return spontaneously over time, a process termed as spontaneous recovery [7, 12]. Thus, extinction is generally thought to result from new inhibitory learning rather than the erasure of an original memory [8, 13, 14], although it has been proposed to involve an unlearning process against the original fear memory [15–17]. Based on the hypothesis that extinction is a new learning and forms an extinction memory, this new memory is more readily forgettable than the original fear memory, i.e., it easily becomes inaccessible and loses the memory control over behavior. The cellular representations that underpin this inherent forgetfulness [18] for the extinction memory remain unclear.

Sparsely distributed populations of neurons activated during specific learning epochs (i.e., memory engram neurons) [19–21] have been increasingly recognized as the cellular representations of memory traces. Memory engram neurons are sufficient and necessary to drive memory retrieval and therefore act as a crucial component of distributed memory traces. Furthermore, memory engram neurons undergo enduring changes in the term of engram neuron-specific connectivity (i.e., engram connectivity) to accommodate the storage and retrieval of memory information [20, 22, 23], although the mechanistic details are unknown. Although it is widely accepted that synaptic plasticity encodes memories, most of previous studies have been conducted at the population level [24, 25] and have not addressed the question of which neurons in specific brain regions undergo plasticity, and site-specific substrates within engram neurons [17, 26] have been largely unexplored. Remarkably, engram neurons and their organization of fear extinction memory are poorly identified [27–29]. Intriguing questions remain regarding where engram neurons of fear extinction memory are assigned and how engram neurons from different brain regions constitute a functional engram circuit and work in concert to store extinction memory.

In this study, we used auditory fear conditioning and extinction paradigms together with an activity-dependent neuronal labeling strategy [30, 31] to identify fear extinction memory engram neurons that constitute directional engram synaptic connectivity to store fear extinction memory and mediate the retrieval in a tripartite neuronal circuit that typically consists of the amygdala, prefrontal cortex, and hippocampus [8]. The engram connectivity is compromised upon spontaneous recovery of the extinguished fear memory, whereas either additional extinction learning or optogenetic induction of long-term potentiation reverses the adaptive shrinkage to prevent the return of fear. We therefore propose that evolvable engram synaptic connectivity between multiple engram neuron populations located in different brain regions collectively constitutes the specific engram construct that governs the retrieval of fear extinction memory.

## Materials and methods

### Animals

All animal procedures were approved by the Animal Ethics Committee of Shanghai Jiao Tong University School of Medicine and by the Institutional Animal Care and Use Committee (Department of Laboratory Animal Science, Shanghai Jiao Tong University School of Medicine; Policy Number DLAS-MP-ANIM. 01–05). The following animals were used in this study: The Fos^2A-iCreER^ (TRAP2) (stock no. 030323) mice were purchased from the Jackson Laboratory (Maine, USA). Prof. Miao He (Fudan University, Shanghai, China) generously gifted the lox-stop-lox-H2B-GFP (H2B-GFP^flox^) reporter mice. All mice were group-housed on a 12-h light/dark cycle with rodent chow and water *ad libitum*. Adult male mice (8–12 weeks old) were used for all experiments.

### Fear conditioning, extinction, and memory test

All auditory fear conditioning, extinction, and memory retrieval procedures were performed using the Ugo Basile Fear Conditioning System (UGO BASILE srl). Full details of fear conditioning, extinction, and memory test are in Supplementary Methods Online.

### Virus constructs

A detailed description of virus constructs is in Supplementary Methods Online.

### Stereotaxic surgery

A detailed description of stereotaxic surgery is in Supplementary Methods Online.

### Engram labeling

Recombination was induced with 4-hydroxytamoxifen (4-OHT, Sigma-Aldrich, Catalog no. H6278). Full details of engram labeling are in Supplementary Methods Online.

### Optogenetic manipulations

For photostimulation during behavioral assays, a 473-nm (blue light) or 638-nm (red light) light-emitting diode (LED) (Hangzhou Newdoon Technology Co. Ltd) was connected to a patch cord with connectors on each end. Full details of optogenetic manipulations are in Supplementary Methods Online.

### Slice electrophysiology

Whole-cell recordings were performed in acute brain slices from mice as described in Supplementary Methods Online.

### *In vivo* optical LTP

Four weeks after extinction learning and engram labelling, mice were placed in homecage after patch cords were fitted to the fiber implants. After a 15 min acclimation period, mice with ChR2^+^ engram terminals in the mPFC received the optical LTP protocol [32] (100 blue light pulses of 2 ms each at a frequency of 100 Hz, repeated 5 times every 3 min). This *in vivo* protocol was repeated 10 times over a 3 h duration. After induction, mice remained in their homecage for an additional 15 min. The behavioral test was then performed one day later.

### Histology and fluorescent immunostaining

A detailed description of histology and fluorescent immunostaining is in Supplementary Methods Online.

### Statistical analyses

A detailed description of the statistical analyses is in Supplementary Methods Online.

## Results

### Retrieval of extinction memory recruits a subset of engram neurons

To identify the engram neurons of fear extinction memory, we trained mice first in auditory fear conditioning and subsequently in extinction learning, during which mice learned to associate auditory cues with footshock first (i.e., fear memory) and then subjected to extinction training with the same auditory cues without footshock (i.e., extinction memory) over 3 days (Supplementary Fig. 1a). After fear conditioning in a particular chamber (context A, Day 1), mice exhibited characteristic freezing behavior as an indicator of associative fear memory for the auditory tone (CS) when presented without footshock (US) in a different test chamber (context B, Day 2–4). Repetitive presentation of the tone alone in context B leads to a gradual decrease in fear response, a process known as extinction learning, resulting in the formation of extinction memory (Supplementary Fig. 1b). Thereafter, reduced freezing response to the tone represented the expression or retrieval of extinction memory. We first characterized the cells activated during retrieval of extinction memory via immunohistochemistry by detecting the protein expression of activity-induced immediate-early gene, c-fos, 90 min after Day 5 of extinction retrieval (Ext. Retr.). Mice in the control group received identical training, although remained in their homecage (HC). Retrieval of extinction memory induced neuronal activation in several brain regions known for extinction memory [8], including medial prefrontal cortex (mPFC), basolateral amygdala (BLA), and ventral hippocampus (vHPC) (Supplementary Fig. 1c, d), supporting these neurons as potential extinction engram neurons.

To map out extinction memory engrams across the whole brain, we used FosTRAP strategy [30, 31, 33] with a reporter line generated by crossing FosCreERT2 (TRAP2) mice with the H2B-GFP (lox-stop-lox-H2B-GFP) reporter line. The targeted recombination in active populations (TRAP) system uses the *Fos* gene locus to drive the expression of tamoxifen-inducible Cre recombinase (CreER), along with a transgenic or virally delivered Cre-dependent effector. When neurons are active in the presence of tamoxifen or 4-hydroxytamoxifen (4-OHT), CreER enters the nucleus to catalyze recombination, resulting in permanent expression of the effector. The TRAP2::H2B-GFP^flox^ (lox-stop-lox-H2B-GFP) double transgenic mice [11] were then trained for fear conditioning and extinction and subsequently injected intraperitoneally with 4-OHT 30 min before Day 4 of extinction, when the fear response was low, resulting in the expression of nuclear histone H2B-binding GFP (H2B-GFP) in neurons activated during extinction (Fig. 1a). We found a large number of brain regions activated by retrieval of extinction memory via whole-brain mapping of H2B-GFP (Fig. 1b, c). Notably, the expression of H2B-GFP in mPFC, BLA, and vHPC coincided with the above c-fos staining induced by extinction memory retrieval (Supplementary Fig. 1), collectively indicating the presence of extinction memory engram neurons in the circuits connected to these brain areas.

**Fig. 1.**
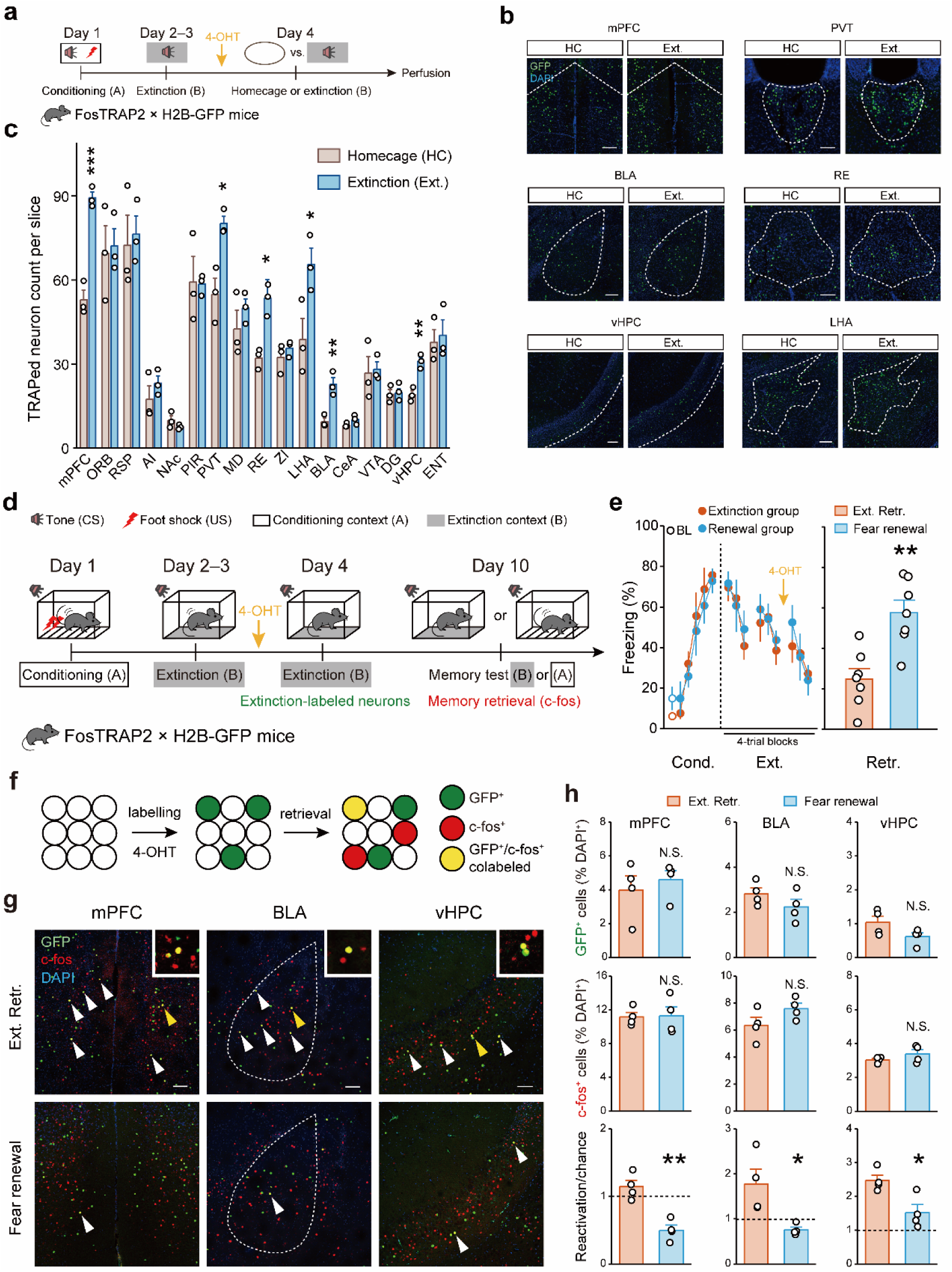
Retrieval of extinction memory recruits a subset of neurons as extinction engram. **a–c** TRAPed neurons activated during extinction. **a** Experimental design. **b** Quantification of TRAPed neurons across 17 brain regions (n = 3 mice per group). *p < 0.05, **p < 0.01, ***p < 0.001, unpaired Student’s *t*-test. mPFC, medial prefrontal cortex; ORB, orbital area; RSP, retrosplenial area; AI, agranular insular area; NAc, nucleus accumbens; PIR, piriform area; PVT, paraventricular nucleus of the thalamus; MD, mediodorsal nucleus of thalamus; RE, nucleus of reuniens; ZI, zona incerta; LHA, lateral hypothalamic area; BLA, basolateral amygdalar nucleus; CeA, central amygdalar nucleus; VTA, ventral tegmental area; DG, dentate gyrus; vHPC, ventral hippocampus; ENT, entorhinal area. **c** Representative images of GFP^+^ cell immunofluorescence. Scale bar, 200 μm. **d** Experimental design for mice subject to fear conditioning (day 1, context A), extinction learning (days 2–4, context B), and subsequent extinction memory test (day 10, context B; referred to as Ext. Retr.) or fear renewal test (day 10, context A; referred to as fear renewal). **e** Freezing responses to the context only (baseline, BL) or CS. While the freezing response during conditioning was calculated by percent freezing time to CS in individual trials, those during extinction learning and memory test were calculated using the average freezing responses of four consecutive trials (4-trial blocks). The same convention was used for the calculation of all behavioral results. Time course of freezing responses to the CS during fear conditioning and extinction learning session (*left*). Statistics are as follows: two-way repeated measures ANOVA, main effect of behavior, conditioning, F_1,72_ = 0.104, p = 0.748; extinction learning, F_1,108_ = 0.539, p = 0.464. Freezing responses to the CS during memory test session (*right*). Data are presented as mean ± S.E.M. **p < 0.01, unpaired Student’s *t*-test. Extinction group, n= 7 mice; renewal group, n = 7 mice. **f** Genetic design to investigate extinction labeled neurons and activated neurons during memory retrieval. The green circles represent labeled neurons during extinction, and administration of 4-OHT to TRAP2 (Fos^2A-iCreER^)::H2B-GFP^flox^ (lox-stop-lox-H2B-GFP) mice was used to induce permanent expression of H2B-GFP in neurons active around the time of the injection. The red circles represent neurons activated during retrieval test. The yellow circles represent colabeled neurons. **g** Representative images of GFP^+^ (*green*) and c-fos^+^ (*red*) immunofluorescence in the mPFC, BLA and vHPC. The white arrowheads denote colabeled GFP^+^/c-fos^+^ cells and the yellow arrowheads denote an enlarged view. Scale bar, 100 μm. **h** Histograms represent mean ± S.E.M. and circles denote individual mice. The extinction group and renewal group displayed similar percentages of GFP^+^ cells among DAPI^+^ cells (*top*). The extinction group and renewal group activated similar percentages of c-fos^+^ cells among DAPI^+^ cells (*middle*). The number of reactivated (GFP^+^/c-fos^+^) cells was significantly higher in the extinction group than the renewal group, the dashed line referred to the value as 1 (*bottom*). Extinction group, n = 4 mice; renewal group, n = 4 mice. N.S., no significant difference, *p < 0.05, **p < 0.01, unpaired Student’s *t*-test.

Reactivation of the cells activated during extinction (GFP^+^ cells) by memory retrieval was assessed by subsequent c-fos staining. The FosTRAP2::H2B-GFP mice received extinction labeling as the presumable extinction engram neurons before Day 4 of extinction. These mice were exposed to an additional extinction memory test or fear renewal test [11] (Fig. 1d). Behaviorally, the renewal test evoked a much higher fear response compared to the extinction test (Fig. 1e). The extinction and renewal test groups displayed comparable numbers of GFP^+^ and c-Fos^+^ cells in mPFC, BLA, and vHPC, as a percentage of 4,6-diamidino-2-phenylindole (DAPI)^+^ cells (Fig. 1f–h). However, consistent with the observation of increased freezing behavior during renewal versus extinction tests, the percentage of GFP^+^/c-Fos^+^ colabeled cells among DAPI^+^ cells divided by the chance percentage [(GFP^+^/DAPI^+^) × (c-fos^+^/DAPI^+^) × 100%] [27, 34, 35] was significantly higher in the extinction than renewal groups (Fig. 1g, h). These results demonstrate that extinction engram neurons are reactivated during extinction retrieval but suppressed by fear renewal. Taken together, these data show that retrieval of extinction memory recruits a unique subset of neurons as the memory engram in the tripartite neuronal circuit consisting of mPFC, BLA, and vHPC.

### Manipulation of extinction engram neurons controls retrieval of extinction memory

Next, we explored the function of extinction engram neurons in the regulation of fear memory-related behaviors. To evaluate whether reactivation of extinction engram neurons is necessary for retrieval of extinction memory, we employed optogenetic inhibition to silence the extinction engram neurons during the extinction test. We injected an adeno-associated virus (AAV) coding for halorhodopsin (NpHR) in a Cre-dependent manner (double-floxed inverted orientated, DIO), along with enhanced green fluorescence protein (EGFP) for visualization (AAV-DIO-NpHR-EGFP), into the mPFC of TRAP2 mice for engram neuronal silencing by illuminating the cell bodies. Mice were bilaterally implanted with optical fibers targeting the mPFC. Three to four weeks later, mice were subjected to fear conditioning followed by extinction learning. The 4-OHT was injected 30 min before the third day of extinction to express NpHR or EGFP in the mPFC extinction engram neurons (Fig. 2a). Six days later, when the effector (i.e., NpHR) was adequately expressed, mice underwent an additional extinction test with optogenetic inhibition. Mice in the NpHR group exhibited a much higher freezing under continuous red light compared to the control group (Fig. 2b), when the firing of extinction memory engram neurons was successfully suppressed by red light (Supplementary Fig. 2). These results indicate that reactivation of extinction engram neurons in mPFC is necessary for retrieval of extinction memory.

**Fig. 2.**
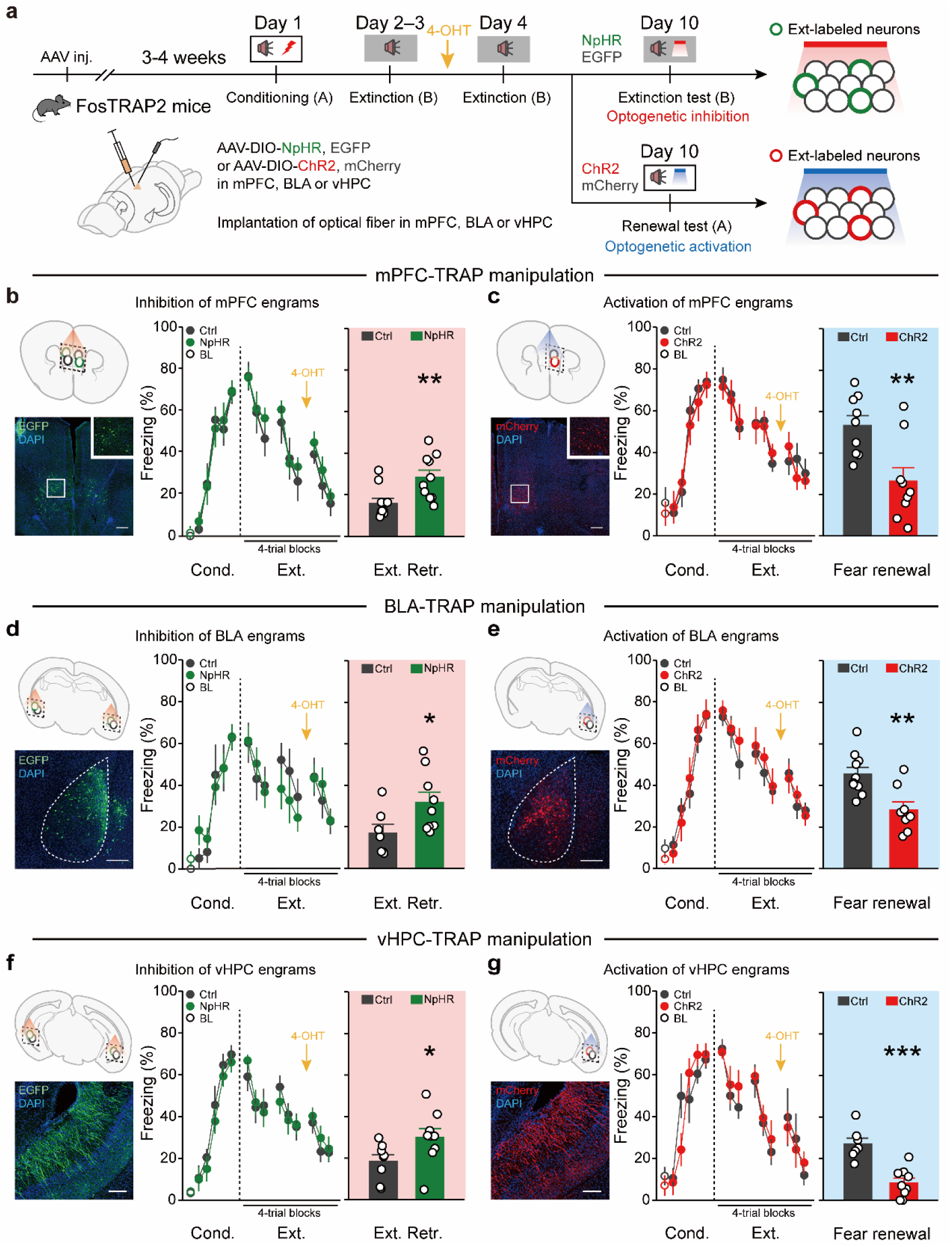
Bidirectional manipulation of extinction engram neurons correspondingly dictates retrieval of extinction memory. **a** Schematics of AAV injections and experimental design. Optogenetic stimulation was delivered during memory test on Day 10. **b, d, f** Loss-of-function studies. Representative images of EGFP expression (*green*) in a mouse that received AAV-DIO-NpHR-EGFP injection into the mPFC, BLA and vHPC (*left*). Scale bar, 200 μm. Time course of freezing responses to the CS during fear conditioning and extinction learning session (*middle*). Statistics are as follows: two-way repeated measures ANOVA, main effect of AAV, (**b**) conditioning, F_1,108_ = 0.046, p = 0.831; extinction learning, F_1,162_ = 2.801, p = 0.096. (**d**) conditioning, F_1,84_ = 0.598, p = 0.441; extinction learning, F_1,126_ = 0.174, p = 0.678. (**f**) conditioning, F_1,90_ = 1.426, p = 0.236; extinction learning, F_1,126_ = 0.150, p = 0.699. Freezing responses to the CS during extinction retrieval session (*right*). Inhibiting extinction engram neurons in mPFC, BLA and vHPC increased freezing levels. Data are presented as mean ± S.E.M. *p < 0.05, **p < 0.01, unpaired Student’s *t*-test. In mPFC group, Ctrl, n = 10 mice, NpHR, n = 10 mice; In BLA group, Ctrl, n = 7 mice, NpHR, n = 9 mice; In vHPC group, Ctrl, n = 8 mice, NpHR, n = 9 mice. **c, e, g** Gain-of-function studies. Representative images of mCherry expression (*red*) in a mouse that received AAV-DIO-ChR2-mCherry injection into the mPFC, BLA and vHPC (*left*). Scale bar, 200 μm. Time course of freezing responses to the CS during fear conditioning and extinction learning session (*middle*). Statistics are as follows: two-way repeated measures ANOVA, main effect of AAV, (**c**) conditioning, F_1,96_ = 0.292, p = 0.590; extinction learning, F_1,144_ = 0.081, p = 0.776. (**e**) conditioning, F_1,96_ = 0.025, p = 0.874; extinction learning, F_1,144_ = 1.435, p = 0.233. (**g**) conditioning, F_1,84_ = 0.123, p = 0.726; extinction learning, F_1,126_ = 0.396, p = 0.530. Freezing responses to the CS during fear renewal session (*right*). Activating extinction engram neurons in mPFC, BLA and vHPC suppressed freezing levels. Data are presented as mean ± S.E.M. *p < 0.05, ***p < 0.001, unpaired Student’s *t*-test. In mPFC group, Ctrl, n = 9 mice, ChR2, n = 9 mice; In BLA group, Ctrl, n = 10 mice, ChR2, n = 8 mice; In vHPC group, Ctrl, n = 7 mice, ChR2, n = 9 mice.

We then determined whether the activation of extinction engram neurons was sufficient to elicit the retrieval of extinction memory under conditions that otherwise do not express extinction memory but rather express fear renewal. To this end, we injected an AAV expressing ChR2-E123T/T159C (ChR2), along with mCherry for visualization (AAV-DIO-ChR2-mCherry) into the mPFC of TRAP2 mice for engram neuronal excitation by illumination of cell bodies (Fig. 2a). Mice were similarly expressed ChR2 or mCherry in the extinction engram neurons of mPFC (Fig. 2a and Supplementary Fig. 2). In the test for fear renewal, control mice displayed higher freezing to the auditory cue compared to mice in the ChR2 group when exposed to the 20-Hz blue light (Fig. 2c). These results indicate that activation of extinction engram neurons in mPFC facilitated the expression of extinction memory to counteract fear renewal.

Likewise, optogenetic manipulations of extinction engram neurons in either BLA or vHPC, similar to that of mPFC, control the retrieval of extinction memory (Fig. 2d–g). Together, these results demonstrate the necessity and sufficiency of extinction engram neurons in any part of tripartite circuits for retrieving extinction memory, implicating that these three brain regions are interlocked to enable the crosstalk of engram neurons in support of retrieval processing.

### Directional connectivity between engram neurons underlies retrieval of extinction memory

Next, we sought to delineate the directionality of mPFC, BLA, and vHPC engram neurons for extinction memory retrieval. In the interconnected hardwired circuits of mPFC-BLA-vHPC, there are bidirectional projections between BLA and mPFC [36–40] and unidirectional projection from vHPC to mPFC [41, 42] involved in extinction memory. We first determined whether there is direct synaptic connectivity between BLA or vHPC neurons and mPFC engram neurons for extinction memory, we applied monosynaptic rabies virus (RV) tracing in TRAP2 mice labeled during extinction. Injection of AAVs with Cre-dependent avian-specific retroviral receptor (TVA) and rabies glycoprotein (RG) into the mPFC followed by injection of RV-expressing EnvA-pseudotyped RG-deleted rabies virus expressing DsRed (RV-ENVA-dG-DsRed) (Fig. 3a), resulted in TVA and RG only in mPFC extinction engram neurons that were labeled with Cre recombinase in an activity-dependent manner, and RV labeling was restricted to presynaptic inputs (Fig. 3b). Trans-synaptically-labeled presynaptic neurons (expressing DsRed only) of mPFC engram neurons were found in multiple cortical and subcortical regions including (most prominently) the piriform area and mediodorsal nucleus of the thalamus, among others (Supplementary Fig. 3). Notably, mPFC engram neurons received inputs from BLA and vHPC, supporting direct connectivity from BLA or vHPC neurons to mPFC engram neurons for extinction memory.

**Fig. 3.**
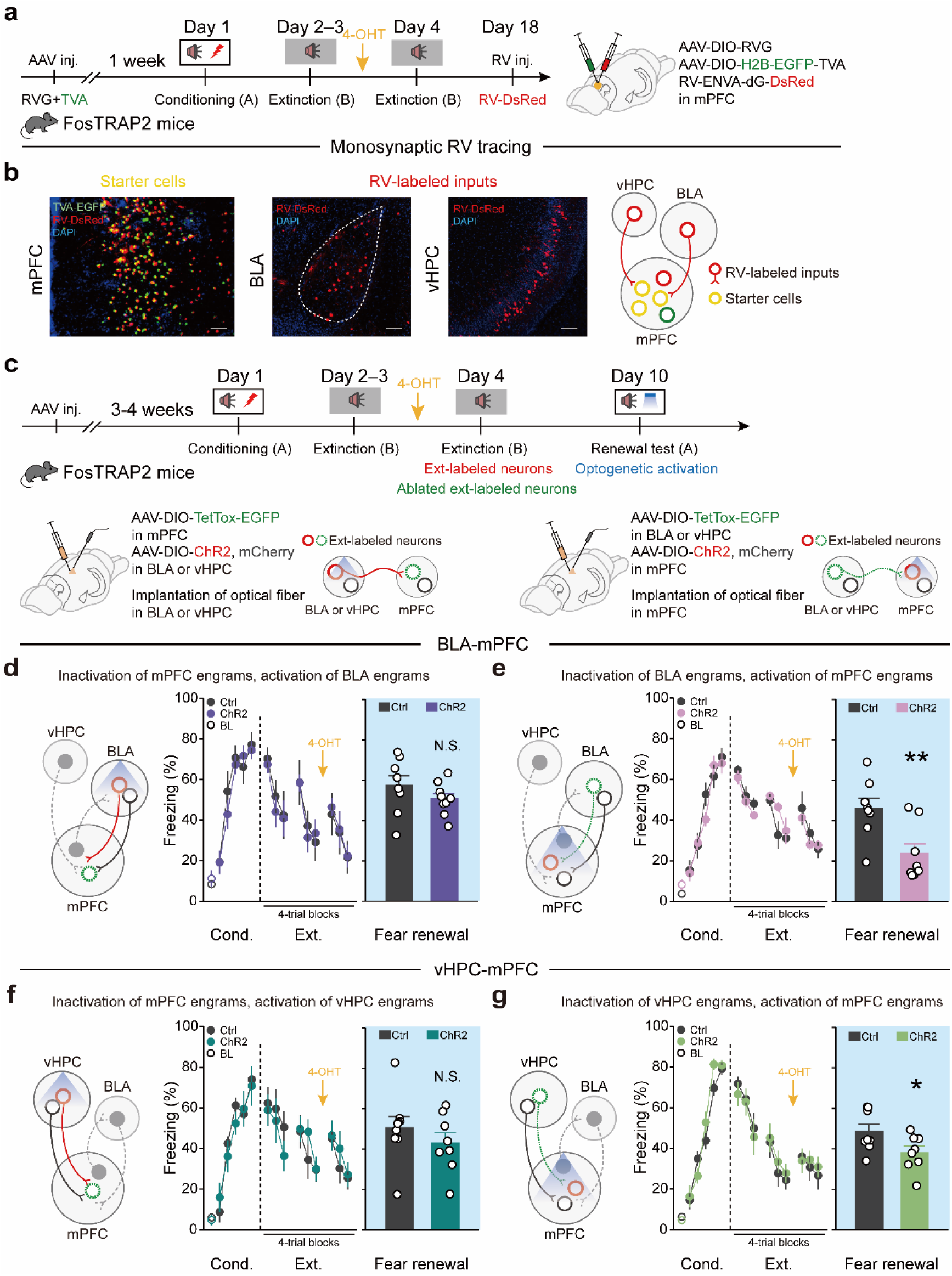
BLA and vHPC→mPFC engram circuits but not the opposite routes are required for extinction retrieval. **a, c** Schematics of AAV injections and experimental design. **b** Representative images of mPFC TVA-EGFP and RV-DsRed injection site and BLA and vHPC DsRed expression. Scale bar, 100 μm. **d–g** Effects of silencing mPFC extinction engram neurons and simultaneously activating BLA or vHPC extinction engram neurons or silencing BLA or vHPC extinction engram neurons and simultaneously stimulating mPFC extinction engram neurons. Experimental diagram (*left*). Time course of freezing responses to the CS during fear conditioning and extinction learning session (*middle*). Statistics are as follows: two-way repeated measures ANOVA, main effect of AAV, (**d**) conditioning, F_1,90_ = 0.208, p = 0.649; extinction learning, F_1,135_ = 0.051, p = 0.822. (**e**) conditioning, F_1,84_ = 0.132, p = 0.718; extinction learning, F_1,126_ = 0.002, p = 0.961. (**f**) conditioning, F_1,90_ = 0.232, p = 0.631; extinction learning, F_1,135_ = 0.022, p = 0.883. (**g**) conditioning, F_1,84_ = 1.154, p = 0.286; extinction learning, F_1,126_ = 0.055, p = 0.816. Freezing responses to the CS during fear renewal session (*right*). Data are presented as mean ± S.E.M. N.S., no significant difference, **p < 0.01, unpaired Student’s *t*-test. In BLA-mPFC group, (**d**) Ctrl, n = 8 mice, ChR2, n = 9 mice; (**e**) Ctrl, n = 8 mice, ChR2, n = 8 mice. In vHPC-mPFC group, (**f**) Ctrl, n = 9 mice, ChR2, n = 8 mice; (**g**) Ctrl, n = 8 mice, ChR2, n = 8 mice.

We then manipulated the synaptic connectivity of engram neurons (i.e., engram connectivity) from BLA or vHPC to mPFC to affect extinction memory retrieval. To silence mPFC engram neurons while activating BLA or vHPC engram neurons, Cre-inducible AAV expressing the light chain of tetanus toxin (TetTox, AAV-DIO-TetTox-EGFP) [43] was injected into mPFC, and Cre-inducible AAV expressing ChR2 (AAV-DIO-ChR2-mCherry) was injected into BLA or vHPC of TRAP2 mice (Fig. 3c). The extinction engram neurons in both mPFC and BLA or vHPC were labeled with different effectors in a single mouse as described previously (see Fig. 2a). We found that silencing mPFC engram neurons led to a failure of optogenetic activation of BLA or vHPC engram neurons to evoke extinction memory retrieval under conditions expressing fear renewal (Fig. 3d, f). Interestingly, in an opposite direction, optogenetic stimulation of mPFC engram neurons still promoted the retrieval of extinction memory despite the silencing of BLA or vHPC engram neurons (Fig. 3e, g). These results suggest that engram connections of BLA→mPFC and vHPC→mPFC, but not that in opposite directions, are necessary for extinction memory retrieval.

### Dynamics of connectivity between engram neurons correlates instability of extinction memory

Notably, extinction memory is not always retrievable and its behavioral impacts on fear memory often subside with passage of time, namely the extinguished fear behavior can return or recover spontaneously over time (i.e., spontaneous recovery). We reasoned that the decline of extinction effects might be due to dynamic remodeling of engram connectivity that leads to inaccessibility of the engram neurons to retrieve extinction memory. To test this hypothesis, we measured adaptive changes of BLA→mPFC and vHPC→mPFC engram connectivity in mice that were tested for spontaneous recovery (S.R.) one month after extinction learning. By contrast, a group of mice received additional extinction learning (Re-Ext.) the day before memory retrieval to re-attain the fear-extinguished state (Fig. 4a–c). We labeled the extinction engram neurons in BLA and vHPC with a blue-light-sensitive opsin, Chronos-EYFP, and red-light-sensitive ChrimsonR-tdTomato, in a counter-balanced manner, and used FosTRAP2::H2B-GFP mice to simultaneously label the extinction engram neurons in mPFC. This procedure allowed for the optogenetic stimulation of BLA→mPFC and vHPC→mPFC engram connectivity separately in mPFC engram or non-engram neurons of the same test subject [44]. By applying paired light pulses to stimulate either BLA or vHPC engram afferents at varying inter-stimulus intervals, we measured the paired-pulse ratios (PPRs) for optically-evoked excitatory postsynaptic currents (oEPSCs), which are inversely related to the presynaptic release probability [45]. In the S.R. group, the PPRs of BLA→mPFC and vHPC→mPFC engram connectivity were both significantly increased in mPFC engram neurons compared with non-engram neurons (Fig. 4d, e), suggesting that spontaneous recovery significantly compromises the presynaptic release probability at BLA→mPFC and vHPC→mPFC engram synapses. In contrast, in the Re-Ext. group, the PPRs of BLA→mPFC and vHPC→mPFC engram connectivity were both significantly decreased in mPFC engram neurons compared to non-engram neurons (Fig. 4d, e). Hence, additional extinction learning re-establishes the lowed setpoint of presynaptic release probability at BLA→mPFC and vHPC→mPFC engram synapses underlying extinction memory.

**Fig. 4.**
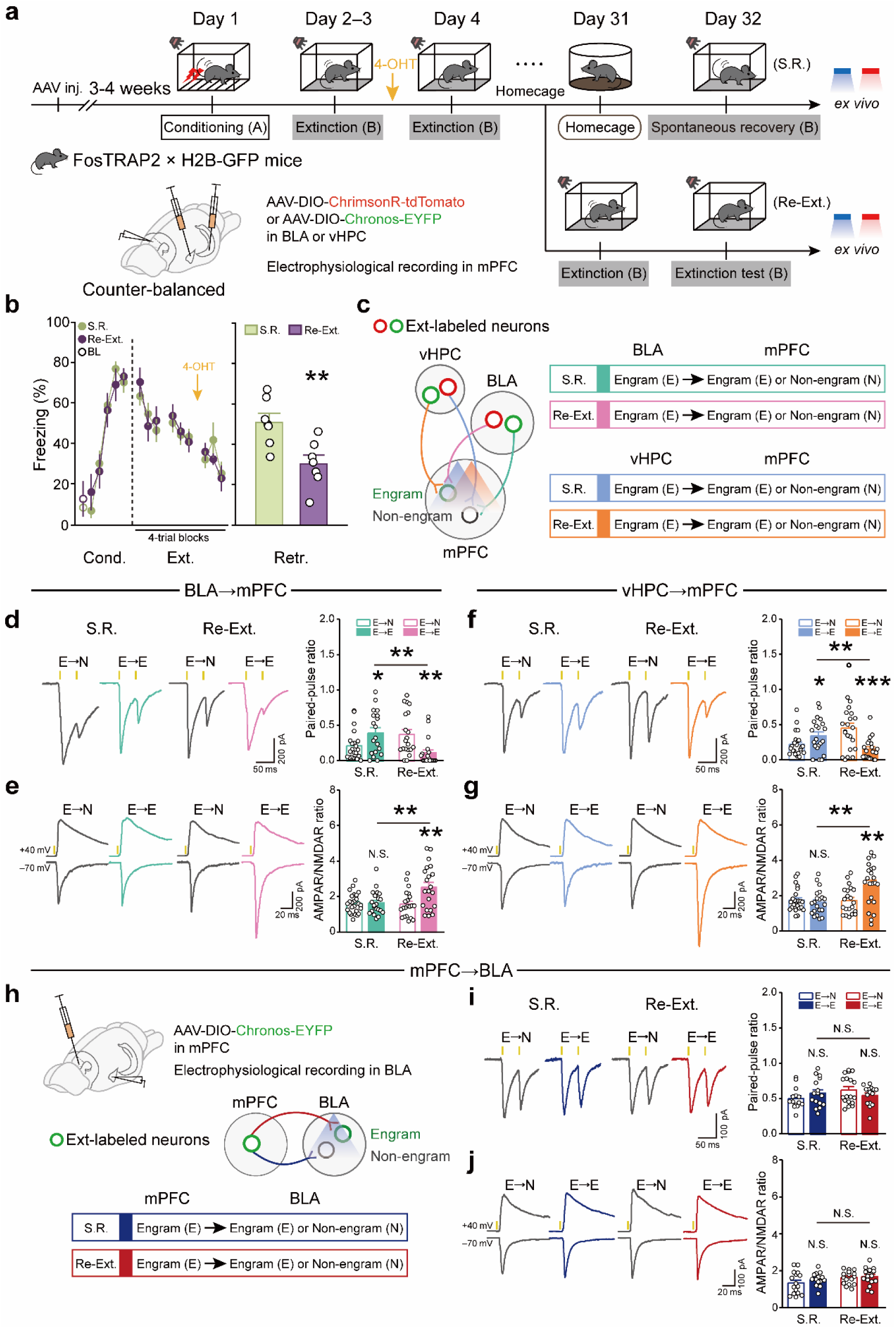
Dynamics of engram connectivity correlate instability of extinction memory. **a** Schematics of AAV injections and experimental design. S.R., spontaneous recovery; Re-Ext., re-extinction before spontaneous recovery. **b** Time course of freezing responses to the CS during fear conditioning and extinction learning session (*left*). Statistics are as follows: two-way repeated measures ANOVA, main effect of behavior, conditioning, F_1,72_ = 0.011, p = 0.918; extinction learning, F_1,108_ = 9.780e-005, p = 0.992. Freezing responses to the CS during memory test session (*right*). Data are presented as mean ± S.E.M. **p < 0.01, unpaired Student’s *t*-test. S.R. group, n = 7 mice; Re-Ext. group, n = 7 mice. **c** Experimental diagram. Schematic diagram of four possible synapse populations among engram and non-engram neurons (*left*). Classification of four synaptic populations indicated by four colors. Turquoise green, E→E or E→N in BLA**→**mPFC pathway during spontaneous recovery; pink, E→E or E→N in BLA**→**mPFC pathway during extinction retrieval; blue, E→E or E→N in vHPC**→**mPFC pathway during spontaneous recovery; dark orange, E→E or E→N in vHPC→mPFC pathway during extinction retrieval (*right*). **d, f** Effects of spontaneous recovery and extinction retrieval on paired-pulse ratios (PPRs) for oEPSCs at the BLA→mPFC (**d**) and vHPC→mPFC (**f**) projections. Representative traces of oEPSCs at the BLA→mPFC (**d**) or vHPC→mPFC (**h**) synapses in the S.R. group and Re-Ext. group induced by paired photostimulations (yellow vertical bars) with a 50-ms interval (*left*). Histograms of mean ± S.E.M. with circles denoting individual neurons (*right*). **d** BLA→mPFC. S.R. group, *p < 0.05, unpaired Student’s *t*-test. E→N, n = 22 neurons from six mice; E→E, n = 19 neurons from six mice. Re-Ext. group, **p < 0.01, unpaired Student’s *t*-test. E→N, n = 20 neurons from six mice; E→E, n = 19 neurons from six mice. **f** vHPC→mPFC. S.R. group, *p < 0.05, unpaired Student’s *t*-test. E→N, n = 23 neurons from six mice; E→E, n = 21 neurons from six mice. Re-Ext. group, ***p < 0.001, unpaired Student’s *t*-test. E→N, n = 22 neurons from six mice; E→E, n = 20 neurons from six mice. **e, g** Effects of spontaneous recovery and extinction retrieval on AMPAR/NMDAR ratios at the BLA→mPFC (**e**) and vHPC→mPFC (**g**) projections. Representative traces of AMPAR- and NMDAR-mediated currents in response to photostimulation of BLA fibers at the BLA→mPFC (**e**) or vHPC fibers at the vHPC→mPFC (**g**) synapses in the S.R. group and Re-Ext. group (*left*). Histograms of mean ± S.E.M. with circles denoting individual neurons (*right*). **e** BLA→mPFC. S.R. group, N.S., no significant difference, unpaired Student’s *t*-test. E→N, n = 25 neurons from six mice; E→E, n = 21 neurons from six mice. Re-Ext. group, **p < 0.01, unpaired Student’s *t*-test. E→N, n = 21 neurons from six mice; E→E, n = 21 neurons from six mice. **g** vHPC→mPFC. S.R. group, N.S., no significant difference, unpaired Student’s *t*-test. E→N, n = 23 neurons from six mice; E→E, n = 21 neurons from six mice. Re-Ext. group, **p < 0.01, unpaired Student’s *t*-test. E→N, n = 21 neurons from six mice; E→E, n = 22 neurons from six mice. **h–j** Adaptations of engram connectivity in mPFC-BLA pathway. **h** Schematics of AAV injections and experimental design. **i** Effects of spontaneous recovery and extinction retrieval on paired-pulse ratios (PPRs) for oEPSCs at the mPFC→BLA projections. Representative traces of oEPSCs at the mPFC→BLA synapses in the S.R. group and Re-Ext. group induced by paired photostimulations (yellow vertical bars) with a 50-ms interval (*left*). Histograms of mean ± S.E.M. with circles denoting individual neurons (*right*). S.R. group, N.S., no significant difference, unpaired Student’s *t*-test. E→N, n = 14 neurons from three mice; E→E, n = 16 neurons from three mice. Re-Ext. group, N.S., no significant difference, unpaired Student’s *t*-test. E→N, n = 18 neurons from three mice; E→E, n = 15 neurons from three mice. **j** Effects of spontaneous recovery and extinction retrieval on AMPAR/NMDAR ratios at the mPFC→BLA projections. Representative traces of AMPAR- and NMDAR-mediated currents in response to photostimulation of mPFC fibers at the mPFC→BLA synapses in the S.R. group and Re-Ext. group (*left*). Histograms of mean ± S.E.M. with circles denoting individual neurons (*right*). S.R. group, N.S., no significant difference, unpaired Student’s *t*-test. E→N, n = 16 neurons from three mice; E→E, n = 14 neurons from three mice. Re-Ext. group, N.S., no significant difference, unpaired Student’s *t*-test. E→N, n = 16 neurons from three mice; E→E, n = 16 neurons from three mice.

Electrophysiologically, we evaluated oEPSCs mediated by glutamatergic α-amino-3-hydroxy-5-methyl-4-isoxazole propionate receptors (AMPARs) and *N*-methyl-D-aspartate receptors (NMDARs), respectively, and noted comparable AMPAR/NMDAR ratios between mPFC engram and non-engram neurons in the S.R. group. However, in the Re-Ext. group, compared with non-engram neurons, the mPFC engram AMPAR/NMDAR ratios were significantly enhanced (Fig. 4f, g), which indicates the postsynaptic adaptions associated with extinction memory instability. As the control, we also measured the adaptive changes mPFC→BLA engram connectivity in both the S.R. and Re-Ext. groups. Similar to before, in the FosTRAP2::H2B-GFP mice, except that extinction engram neurons in mPFC were labeled with ChR2 and that in BLA were simultaneously labeled with GFP (Fig. 4h). Surprisingly, the mPFC→BLA engram connectivity in the term of both presynaptic release probability (Fig. 4i) and postsynaptic receptor activity (Fig. 4j) did not differ between the S.R. and Re-Ext. groups. These results suggest that spontaneous recovery may be due to selective suppression of the directional engram connectivity, and that expression of extinction memory is associated with enhanced strength of engram synapses.

### Optical long-term potentiation of synaptic connectivity between engram neurons enables retrieval of extinction memory

To corroborate the dynamic engram connectivity underlying the instability of extinction memory, we sought to reverse the synaptic strength between BLA or vHPC and mPFC engram neurons using an optical induction approach to rescue extinction memory in lieu of the spontaneous recovery of extinguished fear retrieved by the natural cues [46]. To this end, we utilized optical long-term potentiation (LTP) in specific engram connectivity by activating ultrafast ChR2 mutants with light [47–49]. As before, we selectively expressed ChR2 in BLA and vHPC engram neurons. We performed a high-frequency stimulation (HFS) of ChR2^+^ engram axons in mPFC in the homecage one day before the spontaneous recovery test (Fig. 5a). Remarkably, *in vivo* delivery of engram-specific optical LTP prevented spontaneous recovery of the extinguished fear response and allowed mice to successfully retrieve the extinction memory one month after initial extinction learning (Fig. 5b). The effects of optical LTP were equivalent to those of the re-extinction learning procedure the day before memory retrieval to re-attain the fear-extinguished state (Fig. 5b). Electrophysiologically, spontaneous recovery suppressed the synaptic efficacy of BLA→mPFC and vHPC→mPFC engram connectivity, that PPRs were increased in mPFC engram neurons compared to non-engram neurons -the presynaptic release probability was reduced, but the NMDAR/AMPAR ratios of mPFC engram and non-engram neurons did not differ (Fig. 5c–g), which is consistent with the above findings (see Fig. 4d–g). In contrast, *in vivo* optical LTP protocol abolished the difference in PPRs, but re-extinction learning reversed the changes between mPFC engram and non-engram neurons, and both treatments did re-establish the enhanced mPFC engram AMPAR/NMDAR ratios over non-engram neurons (Fig. 5c–g). These results suggest that dynamic modifications to the synaptic efficacy of engram connectivity are responsible for empowering extinction memory retrieval (Fig. 6), thereby preventing spontaneous recovery of extinguished fear responses.

**Fig. 5.**
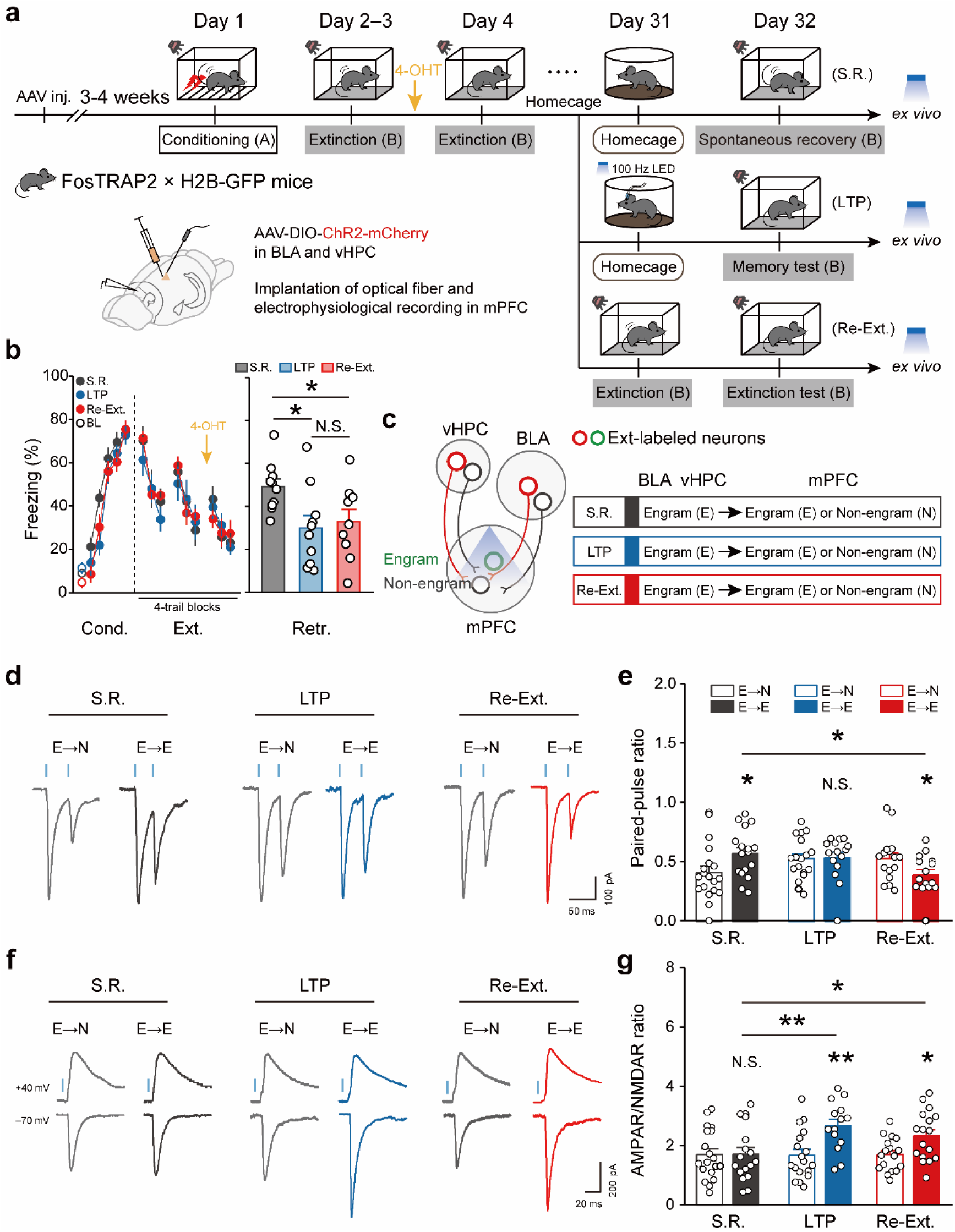
Engram-specific optical long-term potentiation enables retrieval of extinction memory. **a** Schematics of AAV injections and experimental design. S.R., spontaneous recovery; LTP: optical LTP; Re-Ext., re-extinction before spontaneous recovery. **b** Time course of freezing responses to the CS during fear conditioning and extinction learning session (*left*). Statistics are as follows: two-way repeated measures ANOVA, main effect of behavior, conditioning, F_2,156_ = 5.069, p = 0.007; extinction learning, F_2,234_ = 0.347, p = 0.707. Freezing responses to the CS during memory test session (*right*). Data are presented as mean ± S.E.M. N.S., no significant difference, *p < 0.05, unpaired Student’s *t*-test. S.R. group, n = 10 mice; LTP group, n = 10 mice; Re-Ext. group, n = 9 mice. **c** Schematic diagram (*left*). Classification of three synaptic populations indicated by three colors. Dark Grey, E→E or E→N connections during spontaneous recovery; Dark blue, E→E or E→N in connections during optical LTP; red, E→E or E→N in connections during extinction retrieval (*right*). **d** Representative traces of oEPSCs in three groups induced by paired photostimulations (blue vertical bars) with a 50-ms interval. **e** Histograms of mean ± S.E.M. with circles denoting individual neurons. S.R. group, *p < 0.05, unpaired Student’s *t*-test. E→N, n = 19 neurons from six mice; E→E, n = 17 neurons from six mice. LTP group, N.S., no significant difference, unpaired Student’s *t*-test. E→N, n = 18 neurons from six mice; E→E, n = 15 neurons from six mice. Re-Ext. group, *p < 0.05, unpaired Student’s *t*-test. E→N, n = 16 neurons from six mice; E→E, n = 15 neurons from six mice. **f** Representative traces of AMPAR- and NMDAR-mediated currents in response to photostimulation of BLA and vHPC engram fibers at mPFC synapses in three groups. **g** Histograms of mean ± S.E.M. with circles denoting individual neurons. S.R. group, N.S., no significant difference, unpaired Student’s *t*-test. E→N, n = 19 neurons from six mice; E→E, n = 17 neurons from six mice. LTP group, **p < 0.01, unpaired Student’s *t*-test. E→N, n = 18 neurons from six mice; E→E, n = 14 neurons from six mice. Re-Ext. group, *p < 0.05, unpaired Student’s *t*-test. E→N, n = 17 neurons from six mice; E→E, n = 17 neurons from six mice.

**Fig. 6.**
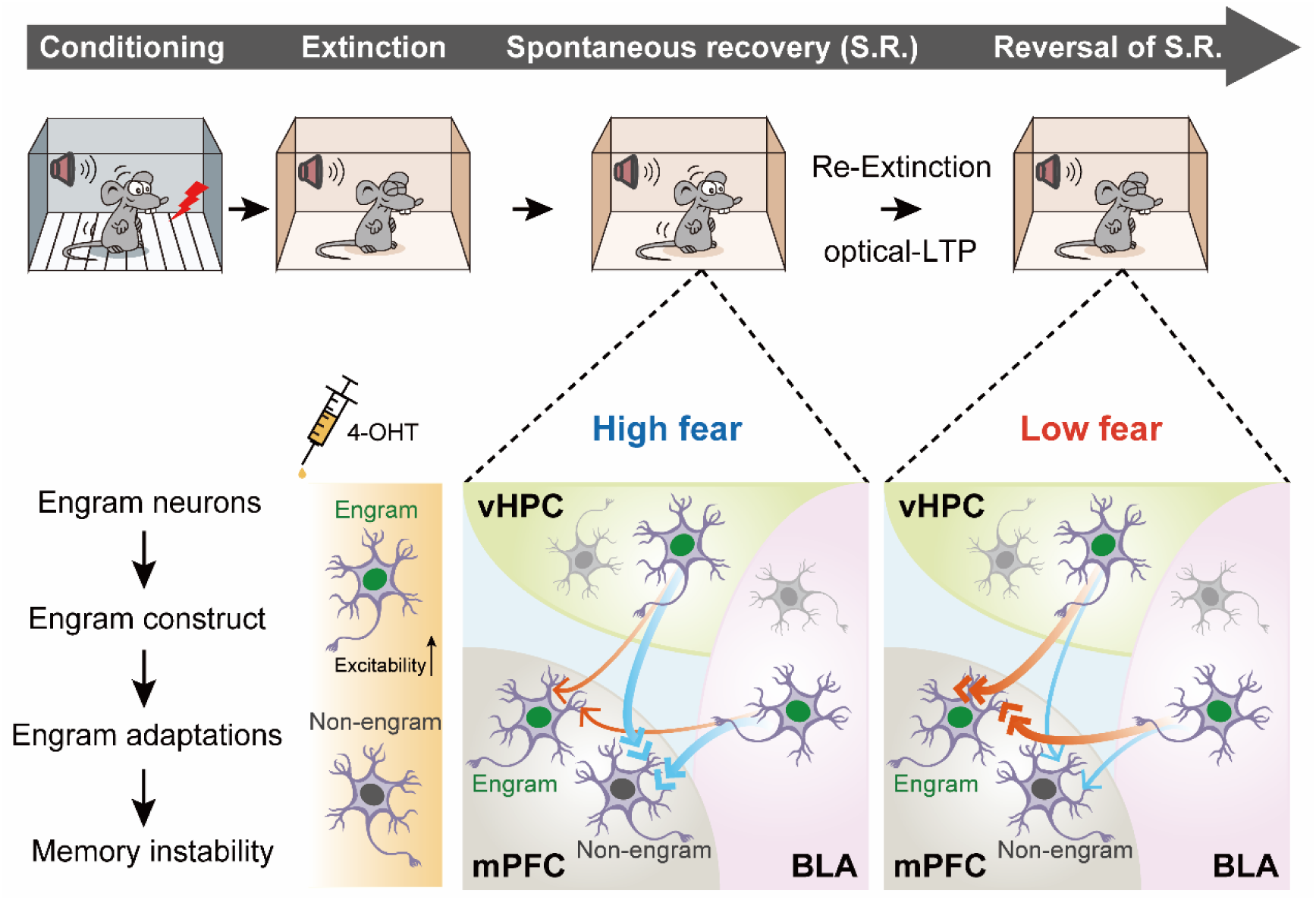
Scheme for a tripartite construct of interregional engram circuits underlies the forgetting of extinction memory. A unidirectional connectivity of new memory engram neurons from BLA and vHPC to mPFC is established during the formation of extinction memory, providing a tripartite construct of circuitry for encoding the extinction memory and storage. Dynamic remodeling of specific engram connectivity dictates both the validity but instability (i.e., readily-forgettable) of fear extinction memory, which enables a longitudinal transformation of memory fate from fear to extinction, relapse and re-extinction. Please see text for more details.

## DISCUSSION

Engram neurons and the cross-region connectivity have been increasingly considered as cellular representations of the memory trace. In this study, we utilized activity-dependent neuronal labeling, manipulation, and recording approaches to dissect the geographic distribution and remodeling of extinction memory engram neurons and their interconnected circuits. Contrary to the “unlearning” hypothesis [17], our results showed that mPFC, BLA, and vHPC serve as the pillar for a tripartite engram circuitry for “new learning” of extinction memory. Our findings established that dynamic evolvability of engram connectivity with bottom-up directionality imparts to the key features of the organization of extinction engram circuits, which ideally explains the readily-forgettable nature of extinction memory. Notably, optogenetic activation of extinction engram connectivity elicited successful retrieval of extinction memory in conditions that would otherwise undergo spontaneous recovery of extinguished fear, suggesting that reinforcement of bottom-up inputs into mPFC from BLA and/or vHPC can counteract relapse of conditioned fear.

The existence of extinction engram neurons in multiple anatomical locations raises an intriguing question of how these distributed cell populations are connected in a manner specific to extinction memory. Here we report that directional synaptic engram inputs from BLA and vHPC to mPFC, but not that with the opposite orientation, constitute a tripartite construct of interregional engram circuits for extinction memory. Moreover, by combining activity-dependent neuronal labeling strategy and monosynaptic RV tracing of the mPFC extinction engram neurons, we anatomically characterized long-range inputs that synapse to mPFC engram neurons, including BLA and vHPC, supporting the direct connectivity from BLA or vHPC neurons to mPFC engram neurons for extinction memory. In addition, the mPFC engram neurons received synaptic inputs from other regions, such as the mediodorsal nucleus of the thalamus (MD), a brain region that is thought to modulate fear extinction [50, 51], and the ventral tegmental area (VTA), supporting the potential involvement of bottom-up prediction error signals in mediating the formation of extinction memory engram [52, 53]. These results support that mPFC engram neurons likely integrate multiple bottom-up signals to form and store extinction memory.

The dynamic changes in synaptic efficacy that occur within different neural circuits have long been thought to be the physical substrate for memory storage and retrieval. In the current study, based on the identification of directional synaptic engram connectivity of BLA→mPFC or vHPC→mPFC for extinction memory, we further characterized a cluster of engram-specific changes in synaptic connectivity strength, including changes in presynaptic transmitter release probability and postsynaptic receptor activities, along with the dynamic and instable fate of extinction memory. Over time, extinction memory engram progressively becomes silent but can be re-vamped to counteract spontaneous recovery of extinguished fear. Indeed, additional extinction learning enables the retrieval of extinction memory by natural clues. We compared the synaptic inputs from BLA or vHPC engram neurons to mPFC engram and non-engram neurons and observed the presynaptic transmitter release probability and postsynaptic AMPAR/NMDAR ratios of mPFC engram neurons increased significantly with extinction than with spontaneous recovery tests, suggesting that specific synaptic potentiation of BLA or vHPC engram is preferentially induced in mPFC engram neurons to achieve the fear-extinguished state. To verify whether this increased plasticity at the mPFC engram neurons represents mnemonic information associated with extinction, we investigated the effect of selectively enhancing postsynaptic AMPAR function in mPFC engram neurons on memory retrieval by optogenetically inducing LTP in the BLA→mPFC and vHPC→mPFC engram connectivity. Our data showed that synaptic potentiation of the BLA→mPFC and vHPC→mPFC engram connectivity, in the homecage, restored extinction memory retrieval via natural auditory cues that otherwise would elicit spontaneous recovery of extinguished fear without additional extinction learning.

In conclusion, we demonstrate that unidirectional connectivity of new memory engram neurons from BLA and vHPC to mPFC is established during the formation of extinction memory, providing a tripartite construct of circuitry for encoding the extinction memory and storage. Dynamic remodeling of specific engram connectivity dictates both the validity but instability (i.e., readily-forgettable) of fear extinction memory. These physically wired engram connectivity for extinction memory storage and retrieval can inform the development of clinical therapies that target excessive traumatic memories associated with numerous neuropsychiatric illnesses, including post-traumatic stress disorder.

## Data availability

All data needed to evaluate the conclusions of the present study are present in the main paper and/or the Supplementary Materials. Additional data are available from the authors upon request.

## Acknowledgements

We thank LetPub (www.letpub.com) for its linguistic assistance and scientific consultation during the preparation of this manuscript. This study was supported by grants from the National Natural Science Foundation of China (31930050, 81961128024, 32071023, 81771214, 31900701, and 81903583), the Science and Technology Commission of Shanghai Municipality (18JC1420302), the Shanghai Municipal Science and Technology Major Project (2018SHZDZX05), the Shanghai Jiao Tong University College of Basic Medical Sciences (YCTSQN2021002), and innovative research team of high-level local universities in Shanghai.

## Supplementary Figures

**Supplementary Fig. 1.**
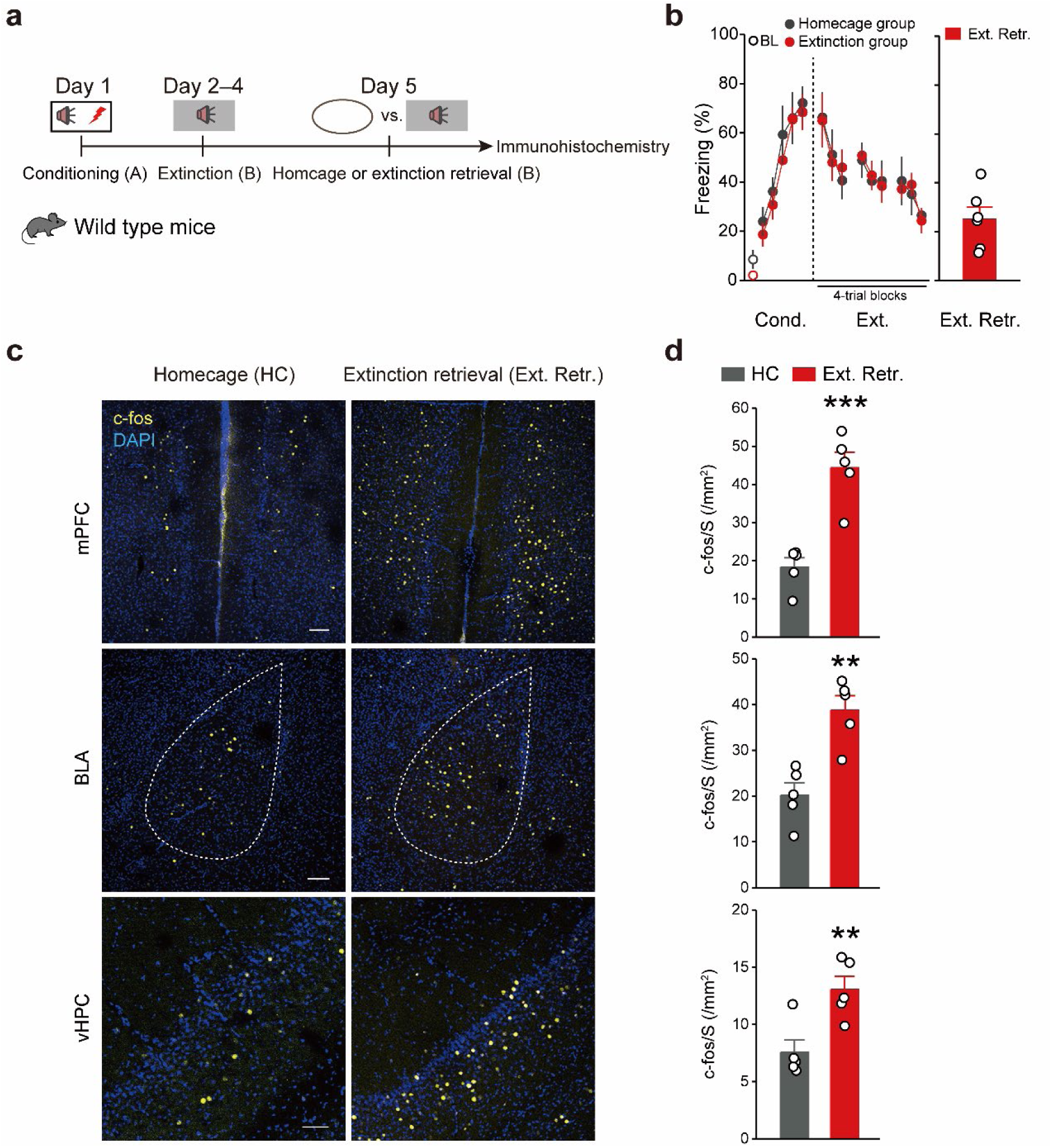
Extinction retrieval activates neurons in mPFC, BLA and vHPC. **a** Experimental design for mice subject to fear conditioning (day 1, context A), extinction learning (day 2–4, context B), and homecage (day 5, homecage; referred to as HC) or extinction retrieval (day 5, context B; referred to as Ext. Retr.). **b** Time course of freezing responses to the CS during fear conditioning, extinction learning and extinction retrieval session. Statistics are as follows: two-way repeated measures ANOVA, main effect of behavior, conditioning, F_1,60_ = 1.953, p = 0.167; extinction learning, F_1,90_ = 0.002, p = 0.962. HC, n = 6 mice; Ext. Retr., n = 6 mice. **c** Representative images of c-fos^+^ (*yellow*) immunofluorescence in the mPFC, BLA and vHPC. Scale bar, 100 μm. **d** The number of c-fos^+^ cells was significantly higher in the Ext. Retr. group than the HC group in the mPFC, BLA and vHPC. (mPFC) HC group, n = 5 mice; Ext. Retr. group, n = 5 mice. (BLA) HC group, n = 5 mice; Ext. Retr. group, n = 5 mice. (vHPC) HC group, n = 5 mice; Ext. Retr. group, n = 5 mice. **p < 0.01, ***p < 0.001, unpaired Student’s *t*-test.

**Supplementary Fig. 2.**
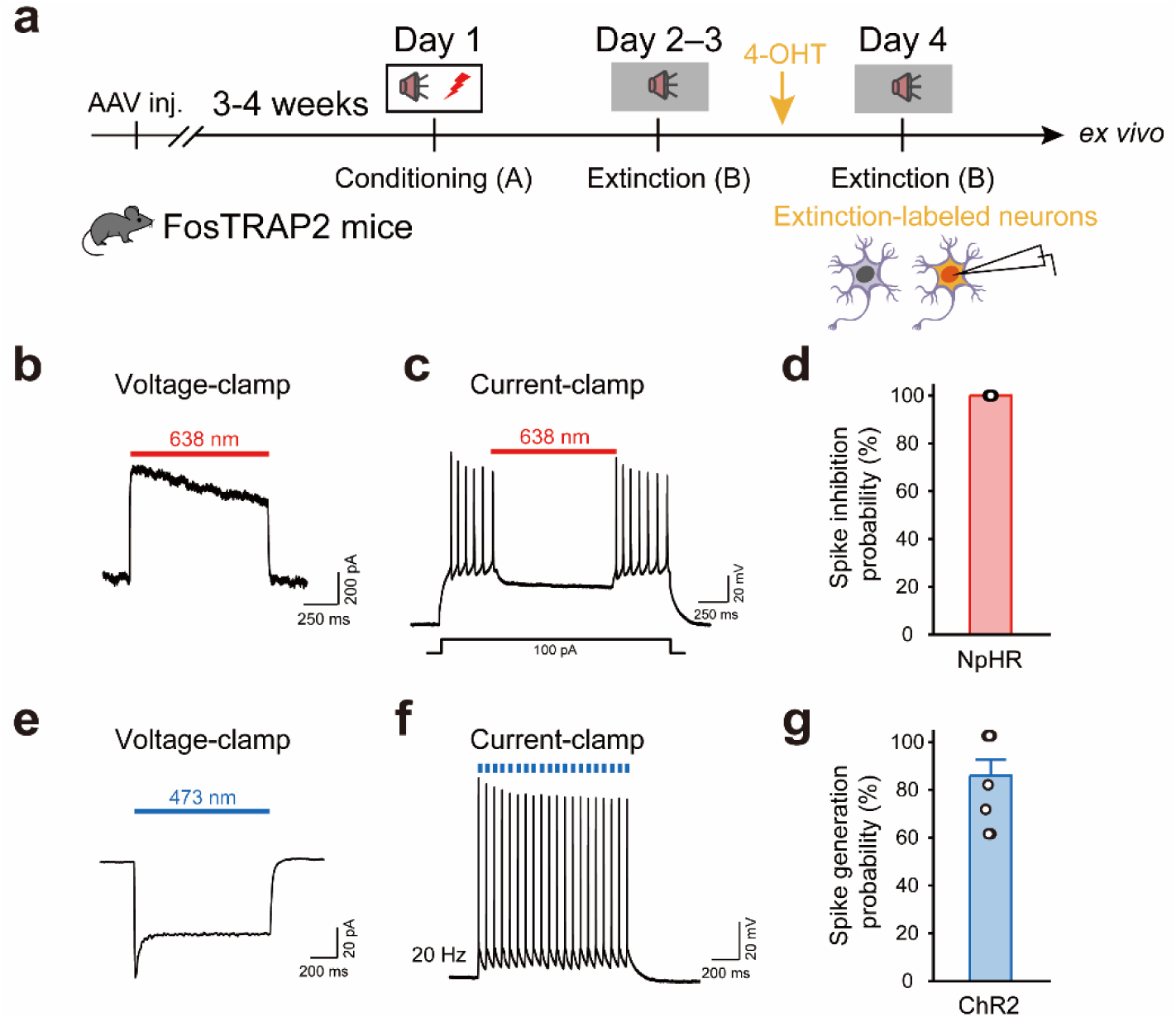
Electrophysiological characterization of optogenetic interventions. **a** Experimental design. **b** Representative trace of red-light-evoked (1 s; continuous) outward current from a NpHR-EGFP-expressing neuron. **c** Blockade of action potential firing recorded from NpHR-EGFP-expressing neurons (100 pA electrical current injection). **d** Bar graph summary of spike inhibition probability under red light. **e** Representative trace of ChR2-mediated current by 1-s pulses of blue light in the voltage-clamp mode. **f** Action potentials were evoked in a ChR2-mCherry-expressing mPFC neuron in current-clamp mode by 1-ms pulses of photostimuli at 20 Hz. **g** Spike generation probability under blue light in ChR2-mCherry-expressing neurons.

**Supplementary Fig. 3.**
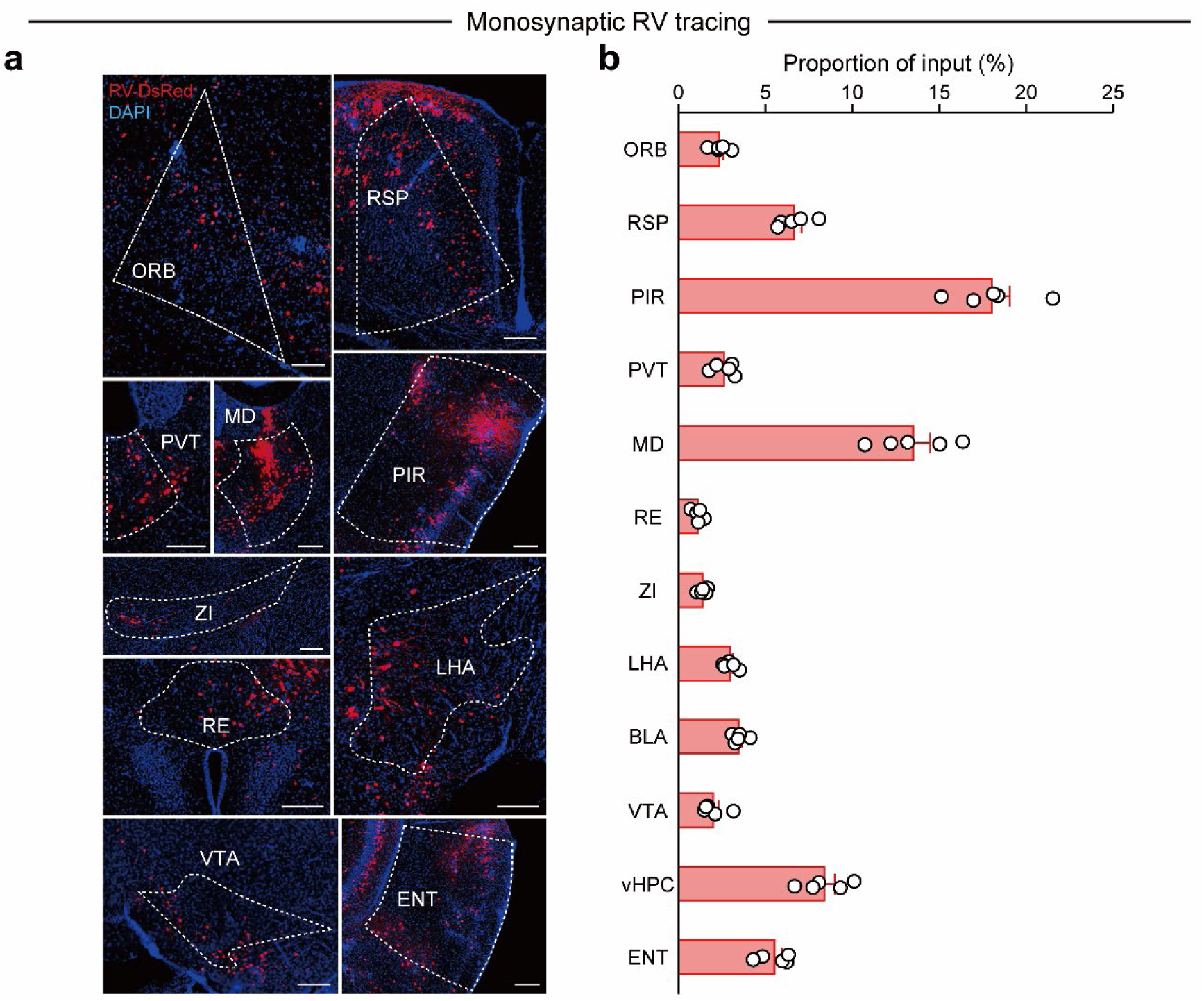
Rabies virus-assisted brain wide mapping of inputs to mPFC extinction engram neurons. **a** Representative images of RV-DsRed expression in major division of the brain. Scale bar, 200 μm. **b** The distribution of RV-DsRed-labeled neurons. ORB, orbital area; RSP, retrosplenial area; PIR, piriform area; PVT, paraventricular nucleus of the thalamus; MD, mediodorsal nucleus of thalamus; RE, nucleus of reuniens; ZI, zona incerta; LHA, lateral hypothalamic area; BLA, basolateral amygdalar nucleus; VTA, ventral tegmental area; vHPC, ventral hippocampus; ENT, entorhinal area. n = 5 mice.

## Supplementary Materials and methods

### Fear conditioning, extinction, and memory test

All auditory fear conditioning, extinction, and memory retrieval procedures were performed using the Ugo Basile Fear Conditioning System (UGO BASILE srl) as described previously [1, 2] with modifications. Briefly, mice were first handled and habituated to the conditioning chamber for five successive days. The conditioning chambers (17 cm × 17 cm × 25 cm) were equipped with stainless-steel shocking grids and connected to a precision-feedback current-regulated shocker (UGO BASILE srl). During fear conditioning, the chamber walls were covered with black-and-white checkered wallpaper, and the chambers were cleaned with 75% ethanol (context A). On day 1, mice were conditioned individually in context A with five pure tones (CS; 4 kHz, 76 dB, 30 s each) delivered at variable intervals (20–180 s). Each tone was co-terminated with a foot shock (US; 0.75 mA, 2 s each). ANY-maze software (Stoelting Co.) was used to automatically control the delivery of tones and foot shocks. Conditioned mice were returned to their home cages 30 s after the end of the last tone, and the floor and walls of the cage were cleaned with 75% ethanol for each mouse.

For extinction learning, mice trained in context A with five CS-US pairings on day 1 were presented with 12 CS presentations (4 kHz, 76 dB, 30 s each) without foot shock in context B (gray floor box) on days 2–4. On day 10, mice received four CS-alone (30 s each) presentations either in the extinction context (context B) for extinction retrieval (also referred to as the extinction test) or in the conditioning context (context A) for fear renewal retrieval (also referred to as the renewal test). Spontaneous recovery tests occurred 28 days after the cessation of extinction learning. During behavioral testing, the chamber was placed in a sound-attenuating enclosure with a ventilation fan and a single house light (UGO BASILE srl). The movement of the mouse in the conditioning or test chamber was recorded using a near-infrared camera and analyzed in real-time with ANY-maze software. A fear response was operationally defined as measurable behavioral freezing (more than 1-s cessation of movement), which was automatically scored and analyzed by ANY-maze software. The time spent freezing during the tone (cue) was measured for each tone presentation.

### Virus constructs

The following viruses were used: AAV-EF1α-DIO-ChR2-E123T/T159C-mCherry (Serotype 2/9), AAV-EF1α-DIO-mCherry (Serotype 2/9), AAV-EF1α-DIO-NpHR3.0-EGFP (Serotype 2/9), AAV-EF1α-DIO-EGFP (Serotype 2/9) were purchased from Obio Technology Co. Ltd. (Shanghai); AAV-EF1α-DIO-TetTox-EGFP and AAV-EF1α-DIO-EGFP (Serotype 2/9) were packaged by Shanghai SunBio Biomedical technology Co. Ltd. (Shanghai); AAV-DIO-ChrimsonR-tdTomato and AAV-DIO-Chronos-EGFP (Serotype 2/9) were produced by Shanghai Taitool Bioscience Co. Ltd. (Shanghai); AAV-EF1α-DIO-RVG (Serotype 2/9), AAV-EF1α-DIO-His-EGFP-TVA (Serotype 2/9), and RV-ENVA-dG-DsRed (titer 3.1 × 10^8^ IFU/ml) were purchased from Brain VTA (Wuhan, China). All viral vectors were stored in aliquots at –80°C until further use. The viral titers for injection were more than 10^12^ viral particles per ml.

### Stereotaxic surgery

Mice at 6–7 weeks old were anesthetized with 1% sodium pentobarbital via a single intraperitoneal injection (10 ml per kg body weight), after which each mouse was mounted in a stereotactic frame with non-rupture ear bars (RWD Life Science). After making an incision to the midline of the scalp, small bilateral craniotomies were performed using a microdrill with 0.5-mm burrs. Glass pipettes, with tip diameters between 10–20 μm were made with a P-97 Micropipette Puller (Sutter Glass pipettes) for AAV microinjections. The microinjection pipettes were first filled with silicone oil and were then connected to a microinjector pump (KDS 310, KD Scientific) with full air exclusion. AAV-containing solutions were loaded into the tips of pipettes and injected at the following coordinates (posterior to Bregma, AP; lateral to the midline, ML; below the Bregma, DV; in mm): mPFC: AP, +1.75 mm; ML, ±0.35 mm; DV, – 2.70 mm; BLA: AP, –1.60 mm; ML, ±3.20 mm; DV, –4.60 mm; vHPC: AP, –3.10 mm; ML, ±3.0 mm; DV, –4.0 mm. After injection, the pipette was left in place for an additional 10 min to allow the injectant to diffuse adequately.

For optogenetic experiments, ceramic fiber optic cannulas (200 μm in diameter, 0.37 numerical aperture (NA), Hangzhou Newdoon Technology) were implanted above the mPFC (AP, +1.75 mm; ML, ±1.80 mm; DV, –2.40 mm; 30° angle); BLA (AP, –1.60 mm; ML, ±3.20 mm; DV, –4.50 mm); vHPC (AP, –3.10 mm; ML, ±3.0 mm; DV, –3.9 mm). The ceramic fiber optic cannulas were positioned in place with acrylic dental cement and secured with skull screws.

### Engram labeling

Recombination was induced with 4-hydroxytamoxifen (4-OHT, Sigma-Aldrich, Catalog no. H6278). In brief, the 4-OHT was dissolved at 20 mg/ml in ethanol by shaking at 37°C for 30 min and was then aliquoted and stored at –20°C for up to several weeks. Before use, 4-OHT was re-dissolved in ethanol by shaking at 37°C for 30 min. Corn oil (Sigma-Aldrich, Catalog no. C8267) was then added for a final concentration of 10 mg/ml 4-OHT, and the ethanol was evaporated by vacuum under centrifugation. The final 10 mg/ml 4-OHT solutions were stored for < 24 h at 4°C before use. All injections were delivered intraperitoneally (i.p.). Mice were transported from the vivarium to an adjacent holding room at least 3 h before the 4-OHT injections to minimize transportation-induced immediate early gene activity. Activity-dependent neuronal labeling was induced by a single intraperitoneal injection of 4-OHT (50 mg/kg mice) administered before the third extinction session for the extinction-labeled mice. Mice were then returned to the vivarium with a regular 12 h light-dark cycle for the remainder of the experiment.

### Optogenetic manipulations

For photostimulation during behavioral assays, a 473-nm (blue light) or 638-nm (red light) light-emitting diode (LED) (Hangzhou Newdoon Technology Co. Ltd) was connected to a patch cord with connectors on each end. The optic fiber implanted in the mouse (200 μm in diameter, 0.37 NA) was connected to the optic patch cord using ceramic mating sleeves. Blue light (473 nm, 4–6 mW) was delivered in 10 ms pulses at 20 Hz during presentation of every 30-s CS (exceeding 5 s before and after the CS to ensure the light delivery covered the CS exposure for the extinction and renewal test). Red light (638 nm) was delivered in a continuous pattern during presentation of every 30-s CS (exceeding 5 s before and after the CS to ensure the light delivery covered the CS exposure for the extinction and renewal test). The final output power ranged from 8–10 mW and depended on the light transmission efficacy of the optical fiber used.

### Slice electrophysiology

Whole-cell recordings were performed in acute brain slices from behaviorally trained mice and/or those that had been stereotaxically injected with AAV-DIO-ChR2-mCherry, or AAV-DIO-NpHR-EGFP or other viruses in different brain regions, as described previously [1, 2] with modifications. Mice were deeply anesthetized with 1% sodium pentobarbital and were subsequently decapitated. Brains were dissected quickly and were chilled in well-oxygenated (95% O_2_/5% CO_2_, v/v) ice-cold artificial cerebrospinal fluid (aCSF) containing the following (in mM): 125 NaCl, 2.5 KCl, 12.5 D-glucose, 1 MgCl_2_, 2 CaCl_2_, 1.25 NaH_2_PO_4_, and 25 NaHCO_3_ (pH 7.35-7.45). Coronal brain slices (300-µm thick) containing regions of interest were cut with a vibratome (Leica VT1000S, Germany). After recovery for 1 h in oxygenated aCSF at 30 ± 1°C, each slice was transferred to a recording chamber and was continuously superfused with oxygenated aCSF at a rate of 1–2 ml per minute. The neurons in mPFC, BLA or vHPC were patched under visual guidance using infrared differential-interference contrast microscopy (BX51WI, Olympus) and an optiMOS camera (QImaging). The slices were continuously perfused with well-oxygenated aCSF at 35 ± 1°C during all electrophysiological studies. Whole-cell patch clamp recordings were performed using an Axon 200B amplifier (Molecular Devices). Membranous currents were sampled and analyzed using a Digidata 1440 interface and a personal computer running Clampex and Clampfit software (Version 10, Axon Instruments). Access resistance was 15–30 MΩ and only cells with a change in access resistance < 20% were included in the analysis. Optical stimulation of ChR2-/ Chronos-, NpHR-, or ChrimsonR-expressing neurons was performed using a collimated LED (Lumen Dynamics) with peak wavelengths of 473, 589, or 638 nm, respectively. The LED was connected to an Axon 200B amplifier to trigger photostimulation. The brain slice in the recording chamber was illuminated through a 40 × water-immersion objective lens (LUMPLFLN 40XW, Olympus). The intensity of photostimulation was directly controlled by the stimulator (2–18 mW/mm^2^), while the duration was set through Digidata 1440 and pClamp 10.5 software. The functional potency of the ChR2-expressing virus was validated by measuring the number of action potentials elicited in neurons of different brain regions using different frequencies of blue-light stimulation (1 ms, 5, 10 and 20 Hz) and the inward photocurrents (1 s pulse) mediated by ChR2 in brain slices. To corroborate the functional potency of NpHR-mediated optogenetic inhibition, red light (λ = 638 nm) was delivered to generate outward photocurrents (1 s pulse) under voltage-clamp mode, promote membrane hyperpolarization, and reduce spikes to current injection under current-clamp mode.

#### Light-evoked EPSCs

To evoke synaptic responses in the mPFC by optogenetic photostimulation of BLA, or vHPC axons, each slice was illuminated every 20 s with blue-or red-light pulses of 5-ms durations according to the light sensitivity of the expressed opsin type (Chronos or ChR2, blue, λ = 473 nm; ChrimsonR, red, λ = 638 nm). To prevent polysynaptic activities from detection in EPSC recordings, we applied appropriate photostimulation intensities that produced 30–50% of the maximal synaptic response. For recording light-evoked EPSCs, the recording pipettes (3–5 MΩ) were filled with a solution containing the following (in mM): 132.5 cesium gluconate, 17.5 CsCl, 2 MgCl_2_, 0.5 EGTA, 10 HEPES, 4 Mg-ATP, and 5 QX-314 chloride (280–300 mOsm, pH 7.2 with CsOH). To determine the paired-pulse ratio (PPR), the patched mPFC neurons were voltage clamped at –70 mV. The AMPAR oEPSCs were evoked by paired photostimulations (50-ms interval; 5-ms duration) of opsin-expressing axons and PPRs were calculated as the peak amplitude ratio of the second to the first oEPSC. To determine the NMDAR/AMPAR ratio, the AMPAR-mediated oEPSCs were recorded at –70 mV while the NMDAR-mediated oEPSCs were quantified as the average EPSC amplitude from 60 to 65 ms after the onset of photostimulation at +40 mV. For each mPFC neuron, photostimulation of the same intensity and duration was used to record the AMPAR-and NMDAR-mediated oEPSCs.

### Histology and fluorescent immunostaining

Animals were deeply anesthetized with 1% sodium pentobarbital and were transcardially perfused with 1 × PBS, followed by 4% paraformaldehyde in 1 × PBS. Brains were extracted and post-fixed overnight at 4°C in 4% paraformaldehyde and then transferred to a 30% sucrose in 1 × PBS medium at 4 °C for 2 d; 30-μm coronal sections were collected using a vibratome (VT1000S, Leica). After a 15-min incubation in 4,6-Diamidino-2-phenylindole dihydrochloride hydrate (DAPI) solution (1: 2000), sections were washed three times (15 min each time) in PBS with 0.1% Tween-20. Slides were mounted in the dark with glass coverslips using mounting media. The coverslips were sealed to the slide with nail polish. All fluorescent images were collected by taking serial z-stack images through 10 × or 20 × objectives of a confocal microscope (Digital Eclipse A1R+, Nikon).

For the c-fos staining, brain slices were washed three times (5 min each time) with 1 × PBS and were then blocked with 10% normal donkey serum in 1 × PBS with 0.3% Triton X-100 (PBST) for 1 h, after which they were incubated overnight at 4°C with rabbit anti-c-fos (1:500, Cell Signaling Technology, catalog no. 2250). Sections were then washed with PBS, incubated in 2% normal donkey serum for 10 min, and then incubated for 2 h with Alexa Fluor® 568 donkey anti-rabbit IgG (H+L) (ThermoFisher Scientific, USA; catalog no. A10042). Sections were washed in 1 × PBS with 0.1% Tween-20, mounted onto slides, and cover slipped with ProLong Gold Antifade Mountant (Invitrogen). Quantification was performed by counting the number of c-fos-positive cells. All counts were performed blind with respect to treatment groups. In Fig. 1h, the amount of reactivation was normalized for chance overlap by dividing the percentage of colabeled^+^ cells among DAPI^+^ by chance [(GFP+/DAPI+) × (c-fos+/DAPI+) × 100%] [2–5].

### Statistical analyses

No statistical methods were used to predetermine sample sizes, but the sample sizes used are similar to those generally employed in the field. The data were collected and processed randomly. The data were collected and processed randomly. All behavioural tests and analysis were blindly conducted. Data distributions were tested for normality and variance equality among groups was assessed using Levene’s test. No data points were excluded. Most histograms display individual data points that represent the values and numbers of individual samples for each condition. Data are presented as the mean ± the standard error of the mean (S.E.M.) unless indicated otherwise. Statistical comparisons were performed using unpaired Student’s *t* tests as well as one-way analyses of variance (ANOVAs) or two-way repeated-measures ANOVAs, where appropriate. For *post-hoc* analysis, we used Bonferroni’s corrections for multiple comparisons. Statistical analysis was performed with IBM SPSS Statistics 25 (SPSS), and p < 0.05 was considered statistically significant. Significance is mainly displayed as *p < 0.05, **p < 0.01, ***p < 0.001; N.S. denotes non-significant values. Not significant values are not denoted except for emphasis.

